# Eukaryotic initiation factor 3F (eIF3F) regulates the IRES-mediated translation of Bcl-xL via its interaction with programmed cell death 4 (PDCD4) protein

**DOI:** 10.1101/2024.03.04.583409

**Authors:** Veda Hegde, Divya K. Sharma, Harshil Patel, Pavan Narasimha, Jason Luddu, Martin Holcik, Nehal Thakor

## Abstract

Programmed cell death 4 (PDCD4) protein is a well-characterized tumor suppressor protein. PDCD4 inhibits mRNA translation by inhibiting the activity of an RNA helicase, eukaryotic initiation factor 4A (eIF4A). We have previously reported that PDCD4 interacts with the internal ribosome entry site (IRES) element that is found within the 5’ untranslated region (UTR) of mRNA encoding B-cell lymphoma extra-large (Bcl-xL) protein. PDCD4’s interaction with the Bcl-xL IRES element inhibits the IRES-mediated translation initiation on Bcl-xL mRNA. However, S6 kinase (S6K)-mediated phosphorylation of PDCD4 activates its degradation by proteasomal degradation pathway and derepress IRES-mediated translation initiation of Bcl-xL mRNA. Interestingly, eIF3F (one of the 13 subunits of eIF3) was reported to recruit S6K to phosphorylate eIF3. Therefore, we were intrigued by the possibility of co-regulation of PDCD4 and eIF3F by S6K and the regulation of IRES-mediated translation initiation by PDCD4-eIF3F. To this end, using co-immunoprecipitation (co-IP), we demonstrated that PDCD4 interacts with several subunits of eIF3. Reciprocal co-IP, endogenous IP, and *in vitro* pull-down assays demonstrated that eIF3F directly interacts with PDCD4 in an RNA-independent manner. In order to functionally characterize the PDCD4-eIF3F complex, we depleted PDCD4 from the glioblastoma (GBM) cells, which resulted in decreased levels of eIF3F. Also, depletion of eIF3F from GBM cells reduced the levels of PDCD4 protein. However, this was not observed in non-cancer cells. Overexpression of PDCD4 resulted in enhanced levels of eIF3F, and *vice versa*. We further confirmed that the interaction of eIF3F and PDCD4 proteins prevents each other’s proteasomal degradation. By performing RNA-IP, we showed that PDCD4 and eIF3F interact with Bcl-xL RNA independently. Moreover, our IRES-bi-cistronic reporter assay and polysome profiling experiments demonstrated that eIF3F regulates IRES-mediated translation of Bcl-xL mRNA, likely via its interaction with PDCD4.

**Significance:** This study uncovers the fundamental mechanism of the internal ribosome entry site (IRES)- mediated translation regulation of B-cell lymphoma extra-large (Bcl-xL) mRNA by programmed cell death 4 (PDCD4) protein, and the eukaryotic initiation factor 3F (eIF3F). Our results show that PDCD4 and eIF3F interact with each other directly and they also interact with Bcl-xL mRNA independently. We show that PDCD4 works via eIF3F to regulate Bcl-xL levels. We also show that the PDCD4-eIF3F-dependent mechanism of Bcl-xL mRNA translation is implicated in glioblastoma (GBM) cells, including the primary brain tumor stem cells (BTSCs), and would likely affect the GBM pathophysiology.

## Introduction

Under normal conditions, most eukaryotic mRNAs utilize the 5’ m7G cap structure of the mRNA to begin protein synthesis (canonical translation initiation) [1–4]. However, during cellular stress conditions, certain eIFs required for canonical translation undergo post-translational modification or proteolytic cleavage, resulting in attenuation of mRNA translation initiation [2, 3, 5]. Distinct subsets of mRNA with specialized structures or sequence elements, such as Internal Ribosome Entry Site (IRES), are selectively translated in response to stress conditions [6, 7]. The translation of mRNAs encoding pro-apoptotic, anti-apoptotic, or stress response proteins is frequently controlled through mechanisms involving IRES-mediated translation initiation under stress conditions [4, 7–9].

A significant body of work indicates that the modulation of IRES-mediated translation can be influenced by IRES trans-acting factors (ITAFs) [7, 10, 11]. ITAFs are known to affect IRES structures of specific mRNA to alter their affinity for ribosomes and translation initiation factors [7, 12, 13]. Each ITAF has the capability to either positively or negatively regulate translation [10]. ITAFs modulate the IRES-mediated translation initiation by interacting with the ribosome or other proteins (such as eIFs), changing the mRNA conformation upon binding, and serving as RNA chaperones. [7, 12, 14, 15]. The levels, post-translational modifications, and availability of eIFs and ITAFs are known to modulate the IRES-mediated translation initiation [7, 8, 16].

PDCD4 is a well-established inhibitor of mRNA translation [17, 18]. As such, PDCD4 binds to eIF4A, which results in the loss of eIF4A’s helicase activity and ultimately inhibition of canonical translation initiation [19, 20]. Additionally, PDCD4 is reported to inhibit canonical mRNA translation by binding to the poly(A) binding protein (PABP) [21] and eIF4G [22]. However, our previous work first time demonstrated that PDCD4 can act as an ITAF [23]. As such, PDCD4 negatively regulates IRES-mediated translation initiation by directly binding to IRES elements of Bcl-xL and X-linked inhibitor of apoptosis protein (XIAP) [19]. The mammalian target of rapamycin (mTOR)/ribosomal protein S6 kinase 1 and 2 (S6K1 and S6K2) axis phosphorylates PDCD4 at serine 67(S67), leading to its proteasomal degradation [23–25]. Although the degradation of PDCD4 enhances the activity of canonical translation initiation, the IRES-mediated translation of XIAP and Bcl-xL mRNAs in HEK293T cells is upregulated. [19, 23, 25].

Arguably, the most complex eukaryotic initiation factor is eIF3 which is comprised of 13 non-identical subunits, each having unique roles in translation [26–29]. eIF3 plays a critical role in canonical translation initiation by bridging the 48S ribosome initiation complex with mRNA via its link with eIF4G [30]. Like PDCD4, mTOR/S6K1/S6K2 axis has been shown to regulate the activity of eIF3 [31, 32] [23–25]. As such, S6K1 physically interacts with eIF3F, and to a lesser extent with eIF3G, which leads to the phosphorylation of eIF3 [32]. This, in turn, enhances the mRNA translation [32]. Further, we and others have shown that eIF3 also regulates the translation initiation of IRES-containing mRNAs such as XIAP and HCV [29, 33]. As both PDCD4 and eIF3 regulate IRES-mediated translation initiation and both are controlled by a common mTOR/S6K axis, we were interested in defining their joint role in IRES-mediated translation of a cellular mRNA. In this study, we show that PDCD4 directly interacts with eIF3F. However, to our surprise, mTOR/S6K signaling did not affect their interaction. We also show that the interaction between PDCD4 and eIF3F plays a critical role in regulating the IRES-mediated translation of Bcl-xL mRNA. Interestingly, PDCD4-eIF3F-dependent regulation of Bcl-xL is found in glioblastoma cells, including brain tumor stem cells (BTSCs). However, non-cancer cells WI-38 (fibroblasts) do not seem to have this mechanism in place. Therefore, it is likely that PDCD4-eIF3F-dependent regulation of Bcl-xL may be specifically relevant to GBM pathophysiology and may impact certain aspects of gliomagenesis.

## Materials and Methods

### Cell lines and constructs

Human glioblastoma cell line U343, U373, BT73MR (BTSCs), as well as HEK293T cells, WI-38, S6K double knockout mouse embryonic fibroblast cells (MEFs), and wild-type MEFs were used for the *cellular* assays. All mammalian cell lines (except BTSCs) were maintained in Dulbecco’s Modified Eagle’s Medium (DMEM) (Hyclone) supplemented with 10% v/v fetal bovine serum (FBS), L-glutamine, and penicillin/streptomycin. BTSCs (BT73MR) were cultured in serum-free media supplemented with epidermal growth factor, fibroblast growth factor, and heparan sulphate in a humidified incubator (37°C, 5% CO2) as described elsewhere [34, 35]. Both MEFs lines were obtained from Dr. Dr. Nahum Sonenberg, Department of Biochemistry, McGill University. The plasmids used to express proteins and reporter constructs to monitor cap-dependent and cap-independent translation are tabulated as shown in (Supplemental Table 1).

### Plasmid purification

A single colony of *E. coli* containing the above-mentioned plasmids from the LB agar was inoculated into a 3 mL LB broth containing 100 μg/ml final concentration of ampicillin. The 3 ml starter culture was incubated in a 37 ⁰C shaker for 10 hours. 1 ml from the starter culture was then inoculated into 200 ml LB broth containing 100 μg/ml final concentration of ampicillin. The 200 ml culture was then incubated overnight at 37 ⁰C with constant shaking at 150 rpm. Following instructions, the plasmid was extracted from the following culture according to the Qiagen plasmid maxiprep kit. The quality of the plasmid purified was analyzed on a 0.8% w/v agarose gel and quantified using the Bio-Drop spectrophotometer.

### Transfection

#### Plasmid transfections

Before transfection, cells were trypsinized and counted using the trypan blue staining. 3 × 10^6^ cells were seeded into a 100-mm tissue culture plate. Forty-eight hours after seeding, cells were transiently transfected with the pcDNA3-PDCD4-FLAG and pcDNA3-FLAG plasmid using Lipofectamine^TM^ 2000 transfection reagent (Thermo Fischer Scientific). The DNA plasmid, transfection reagent, and serum-free opti-MEM media were mixed in proportions per the manufacturer’s protocol and incubated at ambient temperature for 20 minutes. The transfection mix was then added to the 100-mm tissue culture dish containing a monolayer of mammalian cells. The cells were incubated at 37 ^°^C for 48 hours in a CO_2_ incubator before lysis. 3 × 10^5^ cells were seeded into a 6-well tissue culture plate (following reverse transfections of siRNA, see below). Forty-eight hours after seeding, cells were transiently transfected with pcDNA3-FLAG, pcDNA3-PDCD4-FLAG, HA-eIF3F, βGAL-CAT (PBIC) and βGAL – Bcl-xL – IRES – CAT (pBcl-xL) plasmids using Lipofectamine^TM^ 2000 transfection reagent (Thermo Fischer Scientific). The DNA plasmid, transfection reagent, and serum-free opti-MEM media were mixed in proportions per the manufacturer’s protocol and incubated at ambient temperature for 20 minutes. During the incubation period, cells in the 6-well plate were washed with opti-MEM, and 750 μl of opti-MEM was added to each well. 250 µl transfection mix was added to the respective wells. The cells were incubated with the transfection mix for 4-6 hours, and then the opti-MEM media was replaced with DMEM containing 10% v/v FBS. The cells were incubated for 24 hours and harvested.

#### siRNA Reverse transfections

In the case of siRNA transfection, the cells from a 100-mm tissue culture plate were treated with trypsin for 5 min at 37°C. 3 × 10^5^ cells/well were seeded in a 6-well plate. Transfection mix containing Lipofectamine® RNAiMAX (Thermo Fischer Scientific) transfection reagent, siRNA (siPDCD4, sieIF3D, sieIF3E, sieIF3F, sieIF3G – Dharmacon), and opti-MEM media (Thermo Fischer Scientific) were mixed according to manufactures protocol and incubated for 20 minutes at ambient temperature. 250 µl transfection mix was added to respective wells. The plate was set at 37^°^C for 96 hours in a CO_2_ incubator and harvested.

### shRNA-mediated depletion of PDCD4 and eIF3F

Lentiviral particles coding small hairpin (shRNA) specific for PDCD4, eIF3F, scrambled-control was produced by transfecting 3.3 × 10^7^ HEK293T cells in a 100-mm tissue culture plate with 5.4 µg packaging plasmid (psPAX2), 0.6 µg envelope plasmid (VSVG), and 6µg PDCD4/eIF3F shRNA plasmid. The transfection was carried out using the XTREME gene transfection reagent (Millipore sigma) and Opti-MEM media (Thermo Fisher) for 18 hours. Following this, the media was aspirated and replaced with high-Bovine Serum Albumin (BSA, Sigma) growth media (DMEM containing BSA). The media was harvested twice at 24 hours and 48 hours and pooled. The media containing lentiviruses were centrifuged at 1,300 rpm for 2 min at ambient temperature to remove any cell debris. The supernatant was collected, passed through a 0.45 µm filter, aliquoted, and stored at − 80 °C. The aliquot was thawed at room temperature prior to transduction. 1.5 × 10^6^ U343 cells were transduced in a 10 cm tissue culture plate with 1 ml lentivirus added dropwise and 5 µg polybrene. When the cells reached 70% confluency, the media was replaced with puromycin (Sigma Millipore) containing media for selection. At 96 hours after transduction, the cells were washed with PBS and harvested for western blot, and polysome profiling experiments. BTSCs (BT73MR) were dissociated with Accutase (Innovative Cell Technologies, San Diego, USA), and the single cell suspension was obtained. The cells were seeded 250,000 cells/well in a 6-well plate. These cells were transduced with shC, shPDCD4, and sheIF3F lentivirus respectively as described elsewhere [36]. The cells were incubated for 24 hours after which the media was changed and supplemented with puromycin (2 μg/ml) for antibiotic selection. The cells were then harvested after 48 hours and lysates were used for western blot analysis.

### Drug/inhibitor Treatments

U343 cells were reverse transfected with siRNA as mentioned in the section above. In the case of siRNA transfections, the time was optimized to 96 hours before harvesting. The cells were treated with optimized concentrations of 80 nM of AZD2014, and MG132 (1.8 mM) at 24 hrs and 16 hrs before harvesting.

### Immunoprecipitation (IP)

#### FLAG-IP

U343 cells were transfected with pcDNA3-FLAG or pcDNA3-PDCD4-FLAG plasmids. Post 48 hours of transfection, the cells were treated with 3.4% v/v formaldehyde for 10 minutes, followed by 10 minutes of incubation with 0.02 mM glycine to remove the excess formaldehyde. The cells were washed with PBS, and then the cells were lysed in 1.1 ml of RIPA buffer. The lysate was centrifuged at 10,000 g for 10 minutes. 1 ml of the lysate was added to 40 μl of equilibrated ANTI-FLAG® M2 Affinity Gel (Sigma). The lysate was incubated with the affinity gel in the cyclomixer for 4 hours at 4 ⁰C. After incubation, the beads were isolated by centrifuging at 5,000 g for 30 seconds. The matrix was washed with 500 μl of 1X PBS four times. The proteins were eluted by adding 40 μl of 2X concentration Laemmli sample buffer (Bio-Rad) and heating for 10 minutes at 98 ⁰C. The samples were analyzed using the Western blotting technique. 1% of the total cell lysate used for IP was loaded in the SDS-PAGE and co-IP samples as input control.

#### eIF3F IP

40 μl of protein G Dynabeads™ was incubated with 1 μg of eIF3F or PAIP1 (negative control) specific antibody for 20 minutes at room temperature, followed by 1 hour at 4 °C. Unbound antibody was washed with PBS and incubated with cell lysate prepared for FLAG-immunoprecipitation. The lysate and beads were incubated for 4 hours at 4 ⁰C. Post incubation, the beads were separated using a magnetic rack and washed with 500 μl of 1XPBS. Proteins bound to beads were eluted with 40 μl of 2X Laemmli sample buffer and analyzed by Western blotting. 1% of the total lysate volume used for IP was used as input control during Western blot analysis.

#### FLAG-IP to detect RNA-independent protein-protein interaction

Two sets of U343 cells containing a 100-mm tissue culture dish were transfected with the pcDNA3-FLAG or pcDNA3-PDCD4-FLAG plasmids. After 48 hours of transfection, set one was harvested in 1 ml of the RIPA buffer supplemented with 250 μg of RNaseA, and set two was harvested in 1 ml of the RIPA buffer supplemented with 200 units of RNase inhibitor. The lysate was centrifuged at 10,000 g for 10 minutes at 4 ⁰C to separate the cell debris. Then the supernatant was used to perform FLAG-IP similarly as mentioned above.

#### Endogenous IP (Endo IP)

100 μl of protein G Dynabeads™ (Invitrogen) was incubated with PDCD4 specific antibody (1:25 in 0.02 % 1X PBST) or FBS (negative control) for 20 minutes at room temperature, followed by 4 hours at 4 °C. 100 mm culture plates were seeded with U343 and when confluency was reached, they were treated with 10 µl of DMSO and 80 nM of AZD2014. The plates were washed twice with cold 1X PBS and collected using 1ml RIPA buffer with protease and phosphatase inhibitors. Post overnight incubation, the beads were washed with PBS and incubated with U343 cell lysates. The lysate and beads were incubated overnight at 4 ⁰C. The beads were separated using a magnetic rack and washed with 500 μl of cold 1X PBS three times. Proteins bound to beads were treated with 40 μl of 2X Laemmli sample buffer at 95 °C for 10 minutes and analyzed by Western blotting. 3% of the total lysate volume used for IP was used as input control during Western blot analysis.

#### Endogenous RNA IP (Endo RNA IP)

100 μl of protein G Dynabeads™ (Invitrogen) was incubated with PDCD4 and eIF3F specific antibody (1:25 in 0.02 % 1X PBST) or FBS (negative control) for 20 minutes at room temperature, followed by 4 hours at 4 °C. 100 mm culture plates were seeded with U343 and when 80% confluency was reached, the plates were washed twice with cold 1X PBS and collected using 1ml RIPA buffer with protease and phosphatase inhibitors. Post overnight incubation, the beads were washed with PBS and incubated with U343 cell lysates. The lysate and beads were incubated overnight at 4 ⁰C. The beads were separated using a magnetic rack and washed with 500 μl of cold nuclease-free 1X PBS three times. RNA was extracted from the beads using the phenol: chloroform method as explained below. The RNA was analyzed by qPCR and the level of Bcl-xL mRNA expression was measured.

### Western blot assay

The harvested cells were lysed in RIPA (50 mM Tris-HCl pH 7.4, 1 mM EDTA, 150 mM NaCl, 1% v/v NP 40, 0.5% w/v deoxycholic acid, 0.05% w/v SDS, protease inhibitors and phosphatase inhibitors) and centrifuged at 10,000 g for 10 minutes to remove the cell debris. The protein concentration of the supernatant was quantified by the Bradford assay, using the 1X Bradford assay reagent from Bio-Rad, and according to the manufacturer’s protocol. 10-40 μg of total proteins from each lysate were separated on SDS-PAGE. The proteins were transferred onto a nitrocellulose membrane and incubated with 10% w/v fat-free milk in 1X PBST (137 mM NaCl, 2.7 mM KCl, 10 mM Na_2_HPO_4_, 1.8 mM KH_2_PO_4_, and 1% tween 20) for 1 hour followed by incubation with primary antibody overnight at 4 ⁰C. The blots were incubated with a species-specific secondary antibody for one hour and visualized using an Amersham Imager 600. Between primary and secondary antibody probing, blots were washed with 0.1% 1X PBST three times at 15 min intervals. The dilutions of antibodies used were as per the manufacturer’s recommendation.

### Puromycin incorporation assay

U343 cells were reverse transfected with siRNA (siC, siPDCD4, and sieIF3F) as mentioned in the section above. In the case of siRNA transfections, the time was optimized to 96 hours before harvesting. 3 µg/ml of puromycin was added to the media 4 h before harvesting the cells. The cells were harvested and analyzed by western blot as mentioned in the following sections.

### Transformation

*Escherichia coli* (*E. coli*) strains such as DH5α (NEB), BL21 DE3 (NEB), BL21 ROSETTA 2 (DE3), PLYSS (Millipore Sigma), and TOP10 (Thermo Fischer Scientific) were used for transformation. DH5α and TOP10 were used for plasmid preparation, BL21 DE3 for PDCD4 and GST protein purification, and BL21 ROSETTA 2 (DE3) PLYSS for eIF3 subunit protein purification. 50 ng of plasmid DNA was added to the 25 μl aliquot of *E. coli* competent cells in a 1.5 mL tube. Cells were incubated on ice for 45 minutes, followed by a heat shock at 42 ⁰C for 30 seconds. The cells were then placed on ice for 5 minutes and transferred into a culture tube containing 1 ml LB broth. The culture tube was incubated in a 37 ⁰C shaker for 30 minutes. The cells were then streaked onto LB agar plates containing 100 μg/mL final concentration of respective antibiotic and incubated overnight in a 37 °C incubator.

### Recombinant protein purification

A single colony of *E. coli* containing His-PDCD4, GST-eIF3F, or GST bacterial expression construct was inoculated into a 3 ml LB broth (containing 100 μg/ml final concentration of ampicillin) and incubated overnight at 37 ⁰C. 1 ml of the starter culture was inoculated into 200 ml of LB broth containing ampicillin and incubated at 37 ⁰C with constant shaking at 150 rpm. The protein expression was induced with 0.5 mM IPTG at 0.6 OD of the *E. coli* cultures. The culture was incubated for 3 hours after induction and then pelleted by centrifuging at 6,000 g for 30 minutes. The pellet was suspended in 20 ml of lysis buffer (50 mM Tris-HCl pH 8, 1 M NaCl, 5% v/v glycerol, 0.5 mM PMSF, and 1 mM β-mercaptoethanol) and lysed using a French press. The lysate was centrifuged at 16,000 g for 30 minutes, and the supernatant was added to a 500 μl equilibrated Glutathione Sepharose 4 Fast Flow affinity medium (GE) or Ni-NTA agarose (Qiagen). The lysate was incubated with an affinity matrix for 1 hour at 4 ⁰C. The lysate was centrifuged at 1,000 g for 2 minutes, and the supernatant was discarded. The matrix was washed with 20 mL of lysis buffer three times. In the case of His-PDCD4 protein purification, 40 mM of imidazole was added to the lysis buffer during the wash step. GST and tagged protein were eluted using 10 mM glutathione in the lysis buffer. His-PDCD4 was eluted with 500 mM imidazole in the lysis buffer. 20 μl of each eluate was separated using SDS PAGE and visualized by Coomassie staining. The eluates were pooled based on purity and concentrated using an amicon-ultra filtration device (Sigma Millipore) for *in vitro* assays. The Bradford assay quantified purified recombinant proteins.

### In vitro protein pull-down assay

40 μl of Glutathione Sepharose 4 Fast Flow affinity medium was equilibrated with 0.1 M TGEM buffer (20 mM Tris-HCl pH 7.9, 20% v/v glycerol, 1 mM EDTA, 5 mM MgCl2, 0.1% v/v NP 40, 1 mM DTT, 0.2 mM PMSF, 0.1 M NaCl). 500 ng of the quantified bait protein (GST-eIF3F) was diluted to a final volume of 250 μl in 0.1 M TGEM buffer. The diluted sample was added to the equilibrated Glutathione Sepharose 4 Fast Flow affinity medium and incubated at 4 ⁰C for 2 hours. The matrix was then washed once with ice-cold 1 M TGEM buffer (20 mM Tris-HCl pH 7.9, 20% v/v glycerol, 1 mM EDTA, 5 mM MgCl2, 0.1% v/v NP 40, 1 mM DTT, 0.2 mM PMSF, 1 M NaCl) and twice with ice-cold 0.1 M TGMC (20 mM Tris-HCl pH 7.9, 20% v/v glycerol, 5 mM CaCl2, 5 mM MgCl2, 0.1% v/v NP 40, 1 mM DTT, 0.2 mM PMSF, 0.1 M NaCl) buffer to remove the unbound bait protein. 500 ng of PDCD4 was diluted to a final volume of 50 μl in 0.1 M TGMC buffer and added to the matrix coated with the bait protein. The prey protein was incubated with the bait protein at 4 ⁰C for 2 hours. The matrix was washed with 0.1 M TGEM buffer to remove the unbound prey protein. The proteins bound to the matrix were eluted by adding a 2X concentration Laemmli Sample Buffer and heated at 98 °C for 5 minutes. 50 ng of His-PDCD4 and eIF3F was loaded into 10% w/v SDS-PAGE and the pull-down samples as 10% input control. Samples were analyzed by western blot.

### Bicistronic Reporter Assay

The IRES containing bicistronic vector, pβGAL - Bcl-xL - IRES – CAT (pBcl-xL), and the control pβGAL – CAT (PBIC), were used as previously described [19, 23] [23]. The plasmids were transformed using an *E. coli* strain TOP10 (Thermo Scientific). The plasmids were then purified and transfected into U343 cells depleted with PDCD4/eIF3F/eIF3G using siRNAs. Post 48 hours of siRNA transfection, the cells were transfected with 2 µg/ml of PBIC/pBcl-xL using Lipofectamine™ 2000. The cells were then harvested at 96 hours as per the manufacturer’s protocol. The levels of βGAL and CAT were measured according to the protocol of the β-Gal Reporter Gene Assay kit (Roche, cat. # 11758241001) and the CAT ELISA kit (Roche, cat. # 11758241001). The ratios of CAT/βGAL were then analyzed to calculate the relative IRES activity.

### Polysome Profiling

U343 cells were lysed from ten 10 cm plates using cold RNA lysis buffer (1.1 mL basic solution: 0.3 M NaCl, 15 mM MgCl_2_·6H_2_O, 15 mM Tris-HCl, pH 7.4, 110 µL 20% Triton X100, 2 µL SUPERase-In (Thermo Fischer), 5.5 µl of 1% cycloheximide (Sigma-Aldrich)). Equal amounts of cell lysates were loaded on linear sucrose gradients ranging from 10% to 50%. A portion of the cell lysate was kept aside to isolate total RNA and to confirm the depletion of eIF3F or PDCD4 by western blot analysis. 600 µl of cell lysate was loaded onto the sucrose gradients and were centrifuged (rotter SW41) at 39,000 rpm for 90 min at 4 °C. Gradients were then fractionated using a density gradient fractionation system (Brandel #SYN-202) at 1 ml each. Polysome profile was read using DATAQ software. RNA from each fraction was isolated and RT-qPCR was performed as explained below.

### RNA isolation

RNA isolation was done using the phenol-chloroform extraction method. 1ml of polysome fraction or whole cell lysate was incubated with 50 µl of Proteinase K solution (57 µl 10% SDS, 7.5 µl 0.5 M EDTA, 1 µl Glycoblue, 4 µl of 20 mg/ml Proteinase K) at 55 ⁰C for 1 hr. Equal volumes of phenol:chloroform: isoamyl alcohol mixture (125:24:1, acidic pH 4.5) was added to each fraction/lysate. The fractions were centrifuged at 15,000 rpm for 5 min at room temperature. The aqueous phase was carefully removed, and an equal volume of chloroform was added. Centrifugation was repeated as mentioned above. To the aqueous phase, 1:10 volumes of 3 M sodium acetate (pH 5.2) and 1.5 volumes of chilled absolute ethanol were added and vortexed for ∼ 15 sec. The tubes were left overnight at −20 ⁰C to precipitate. Overnight incubation was followed by centrifugation at 15,000 rpm for 30 min at 4 ⁰C. The pellets were washed with chilled RNase-free 70% ethanol. Centrifugation was repeated as mentioned above. The pellets were air-dried and resuspended in RNase-free water. The quality of RNA was checked with a nanodrop (A260/280 ∼ 1.7 – 2) and was further used for RT-qPCR analysis.

### RT-qPCR

RNA was isolated as mentioned above and cDNA was generated from equal quantities of RNA using the qScript cDNA synthesis kit (Quanta Biosciences). Quantitative PCR was performed in a CFX-96 real-time thermocycler (Bio-Rad) with PerfeCTa SYBR Green SuperMix (Quanta Biosciences) according to the manufacturer’s instructions. Negative controls without template DNA (NTR) were run in duplicate. Each reaction was run in duplicate with the following cycle conditions: 1 cycle at 95°C for 3 min followed by 35 cycles at 95 °C for 15 seconds, the annealing temperature of 55 °C for 35 seconds, and 72 °C for 1 min. A melting curve step was added to check the purity of the PCR product. This step consisted of a ramp of the temperature from 65 ⁰C to 95°C at an increment of 0.5 °C and a hold for 5 seconds at each step. Expression levels of, Bcl-xL, PDCD4, and eIF3F mRNA’s (relative to β-actin mRNA) were determined using the ΔΔC_t_ method. PDCD4, eIF3F, Bcl-xL, and Actin primers were obtained from Quantitect (Qiagen).

## Results

### eIF3F and PDCD4 interact with each other directly in an RNA-independent manner

As both PDCD4 and eIF3 are regulated by the mTOR/S6K1/2 axis, our preliminary hypothesis aimed at elucidating the role of S6K1/2 in the interaction between PDCD4 and eIF3-subunits. To this end, we first determined if PDCD4 and eIF3 interact with each other. To check the interaction between PDCD4 and eIF3 subunits, we over-expressed FLAG-tagged PDCD4 in U343 glioblastoma cells and performed co-immunoprecipitation (co-IP) using FLAG-specific antibody-conjugated beads. By performing western blot analysis, we confirmed that several subunits of eIF3 (namely eIF3F, eIF3G, eIF3B, eIF3D, and eIF3E) interact with FLAG-PDCD4 (Fig. 1A). Notably, the intensity of the eIF3H band in the FLAG-PDCD4 IP was comparable to that in the FLAG-tag IP control, suggesting non-specific interaction with either the FLAG tag or the affinity matrix. (Fig. 1A). Subsequent probing with anti-eIF3H mouse-raised antibodies revealed a slightly higher molecular weight protein alongside eIF3H, corresponding to the molecular weight of mouse IgG from the anti-FLAG antibody (Fig. 1A). As the Poly(A)-binding protein (PABP) is known to interact with PDCD4, PABP served as a positive control. Indeed, PABP interacts specifically with FLAG-PDCD4, but not with FLAG-tag only (Fig. 1A). Given the known regulatory role of S6K1 in PAIP1 interaction with eIF3 [32], the interaction between PAIP1 and FLAG-PDCD4 was checked. However, PAIP1 did not co-immunoprecipitate with either FLAG-PDCD4 or the FLAG-tag, indicating no interaction (Fig. 1A).

**Figure 1:**
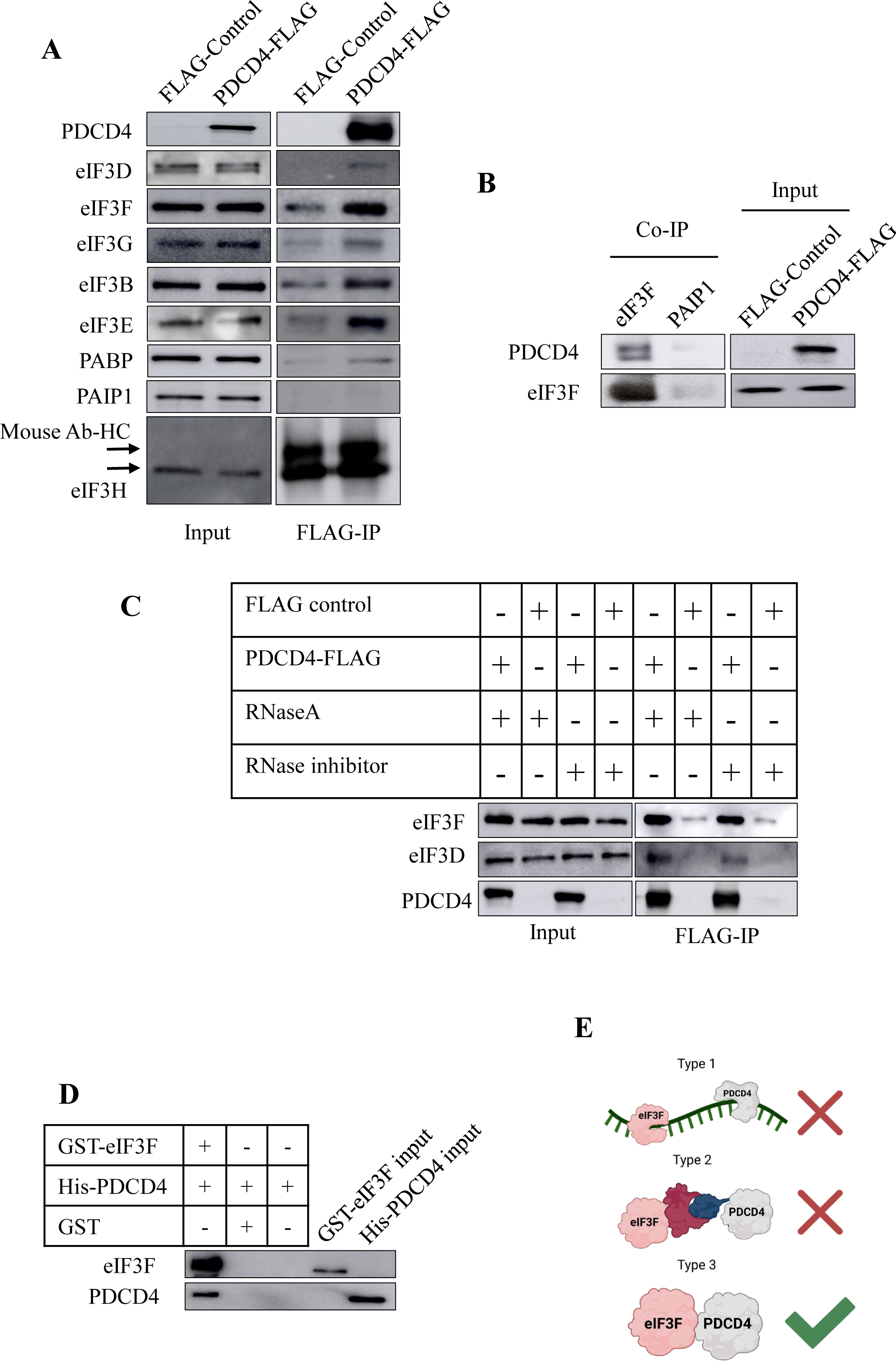
eIF3F and PDCD4 interact in an RNA-independent manner. co-IP samples were probed with the same antibodies as the inputs. eIF3F, eIF3G, eIF3B, eIF3D, eIF3E and PABP were co-immunoprecipitated with FLAG-PDCD4. The result shows that respective eIF3 subunits were immunoprecipitated along with PDCD4 **(A)**. *Reciprocal IP* was performed using eIF3F antibody co-immunoprecipitated FLAG-PDCD4 along with eIF3F as seen in the IP lane **(B)**. PDCD4 was co-immunoprecipitated with eIF3F and eIF3D (right panel, lanes 1 and 3). There was no difference in their interaction between RNaseA treatment and RNAase inhibitor treatment. This indicates that the interaction between PDCD4 and eIF3 is not dependent on RNA **(C)**. The prey protein recombinant PDCD4 binds directly to the bait protein recombinant eIF3F. PDCD4 is pulled down with eIF3F (right column). This suggests direct interaction with each other **(D)**. *Possible types of interactions:* PDCD4 and eIF3F may interact with the same RNA molecule, PDCD4 and eIF3F may be part of the same protein complex, or there could be direct interaction between PDCD4 and eIF3F. **(E)**. Fig. 1E was created by VH using BioRender^TM^.

To further validate the PDCD4-eIF3 interaction, a reciprocal IP was conducted using eIF3F antibodies. Subsequent probing with a conformation-specific secondary anti-rabbit antibody raised in mice confirmed co-immunoprecipitation of PDCD4 with the eIF3F subunit (Fig. 1B). Interestingly, an additional band, potentially representing the FLAG-tagged or post-translationally modified form of PDCD4, was observed along with PDCD4 in the eIF3F immunoprecipitated sample (Fig. 1B). Notably, PDCD4 was not detected in lanes with elution from beads coated with control antibodies (PAIP1) or lysates lacking FLAG-PDCD4 expression. This collective data strongly supports the hypothesis of a specific interaction between PDCD4 and the eIF3F subunit (Fig. 1B). Additionally, the Endo-IP performed using PDCD4 antibody co-immunoprecipitated eIF3F (Fig. 2E; details below).

**Figure 2:**
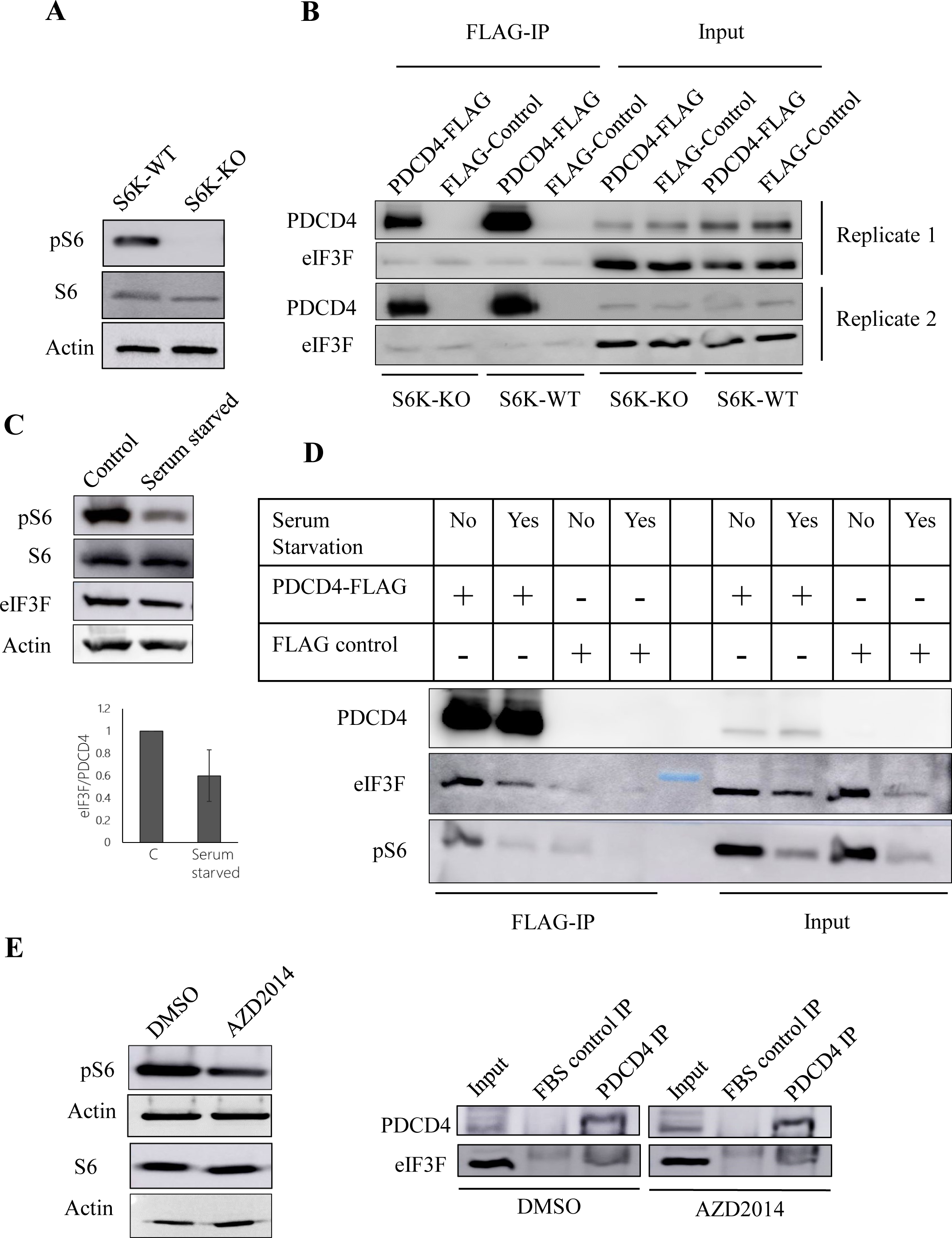
mTOR/S6K axis does not affect eIF3F-PDCD4 interaction. S6 protein was equally expressed in wild-type and S6K double knockout MEFs. However, phosphorylation of S6 is not detected in S6K double knockout MEFs, which suggests the lack of S6K activity **(A)**. PDCD4 and eIF3F protein levels were similar in all four input lanes. In the co-IP lanes, eIF3F band intensity was similar in all four lanes, indicating no difference in interaction with PDCD4. This suggests that PDCD4 and eIF3 interaction is not affected in MEFs by the S6K activity **(B)**. Serum starvation decreased the S6 phosphorylation, which suggests decreased S6K activity **(C)**. eIF3F and pS6 were co-immunoprecipitated with PDCD4. This interaction decreased with serum starvation seen in the co-IP lanes. However, the input lanes indicated a decrease in eIF3F and pS6 protein levels in the total cell lysate. This suggests that the PDCD4-eIF3F interaction is not affected by serum starvation **(D)**. U343 cells were treated with DMSO and a sub-lethal dose of mTOR inhibitor AZD2014. AZD2014 inhibitor activity was confirmed by observing the levels of S6 and pS6 levels. eIF3F is co-immunoprecipitated along with PDCD4 endogenously. However, AZD2014 treatment did not affect endogenous PDCD4-eIF3F interaction **(E)**.

The interaction of eIF3F and PDCD4 proteins demonstrated by co-IP can be a result of three major types of protein association in cells (Fig. 1E). In type 1PDCD4 and eIF3 may interact with the same RNA molecule without interfering with each other’s interactions. In Type 2, it is possible that PDCD4 and eIF3F may participate in a shared protein complex,including additional proteins, consequently leading to their co-immunoprecipitation. This implies the likelihood of a physical association between PDCD4 and eIF3F within the context of a common protein complex. Lastly, in Type 3 PDCD4 and eIF3 subunits can physically interact with each other directly (Fig. 1E).

To check the involvement of the RNA in PDCD4-eIF3 interaction, two separate FLAG IP experiments were performed in the presence of RNAse A enzyme (use of RNAse inhibitor was avoided) and in the presence of RNAse inhibitor (RNAse A enzyme was not used). Both eIF3F and eIF3D were co-immunoprecipitated with FLAG-PDCD4 in both RNase A treated and RNase inhibited cell lysates (Fig. 1C). The protein levels of eIF3F and eIF3D were found to be similar in all four lysates as seen in the input blot. PDCD4 protein was not detected in the FLAG tag control lysate. Additionally, there was no significant interaction of eIF3 subunits with FLAG-tag or the matrix used for IP. Therefore, we inferred that the PDCD4-eIF3F and PDCD4-eIF3D interaction is RNA-independent (Fig. 1C) To investigate if PDCD4 interacts directly with eIF3F, *in vitro* protein pull-down was performed using Glutathione Sepharose Matrix. Western blot analysis of the pulled-down proteins with anti-eIF3F and anti-PDCD4 antibody showed interaction between PDCD4 and eIF3F (Fig. 1D). His-PDCD4 was detected in the presence of GST-eIF3F but not with GST or Glutathione Sepharose Matrix only (Fig. 1D) suggesting a direct protein-protein interaction between eIF3F and PDCD4 (Fig. 1D).

### Interaction between PDCD4 and eIF3F was not affected by S6K activity

S6K1 is known to interact with eIF3F and to regulate the activity of eIF3 [32]. Moreover, both S6K1 and S6K2 are reported to regulate the levels of PDCD4 via the proteasomal degradation pathway [37]. Therefore, we wanted to investigate if S6K1 & 2 play any role in the interaction between eIF3F and PDCD4. To this end, we employed S6K double knockout mouse embryonic fibroblasts (MEFs). First, we performed western blot analysis of phosphorylated form of S6 protein in wildtype and S6K double knockout cells (Fig. 2A). The complete lack of phospho-S6 (pS6) confirms that S6K1 & 2 activity was absent in the S6K double knockout cells (Fig. 2A). Second, to check if PDCD4 and eIF3F interact in the S6K1 and 2 double knockout conditions, FLAG-PDCD4 was expressed in S6K1 & 2 expressing MEFs (wildtype) and S6K1 & 2 double knockout MEFs (Fig. 2B). Subsequently, we performed FLAG Co-IP followed by western blot analysis. The membrane was probed with anti-PDCD4, anti-eIF3F, anti-S6, and anti-pS6 antibodies. There were no significant changes in the level of co-immunoprecipitated eIF3F in FLAG-PDCD4 and FLAG-tag expressed MEFs (Fig. 2B). Therefore, S6K1 & 2 double knockout and wildtype MEFs may not be suitable to test the effect of S6K1 & 2 on the PDCD4-eIF3 interaction as there was no PDCD4-eIF3F interaction detected in these cells. Therefore, we investigated the role of S6K1/2 in PDCD4-eIF3F interaction in U343 cells. To this end, we serum-deprived U343 cells for 24 hours and performed western blot analysis to check the levels of pS6. A reduced phosphorylation of S6 was observed in U343 cells upon serum starvation (Fig. 2C, 2D). The expression of S6 protein remained unaffected during serum starvation (Fig. 2C). As expected, these observations suggest that serum deprivation inhibits the activity of S6K1/2 in U343 cells (Fig. 2C). Subsequently, we expressed FLAG-PDCD4 in U343 cells and performed FLAG Co-IP experiments in control and serum-deprived U343 cells (Fig. 2D). eIF3F and pS6 were found to be co-immunoprecipitated with FLAG-PDCD4 (Fig. 2D). The eIF3F over PDCD4 ratio was found to be decreased by about 25% in serum-starved condition when normalized with the amount of PDCD4 immunoprecipitated (Fig. 2C). However, there was also a similar decrease in the eIF3F levels upon serum starvation in input (cell lysate) lanes (Fig. 2D). The observed reduction in PDCD4-eIF3F interaction may be attributed to a decreased eIF3F protein level rather than a variation in the PDCD4-eIF3F interaction (Fig. 2D). Similarly, the reduction in the PDCD4-pS6 interaction was noted as a likely outcome of reduced S6 phosphorylation during serum starvation (Fig. 2D).

The influence of growth factors on mTOR inhibition is important for the activation of S6K1/2. Accordingly, we treated U343 cells with a sub-lethal dose of mTOR inhibitor (AZD2014) and monitored the phosphorylation of S6 protein. AZD2014 treatment decreased the phosphorylation of S6 protein but did not reduce the levels of S6 protein (Fig. 2E; left panel). We performed an Endo-IP experiment in DMSO (control)- and AZD2014-treated U343 cells using antibodies against PDCD4 (Fig. 2E; right panel). FBS was used as a negative control for IP. Endogenous eIF3F was co-immunoprecipitated with endogenous PDCD4 under DMSO (control) treatment conditions. This endogenous interaction was not affected by AZD2014 treatment (Fig. 2E; right panel). Collectively, these data (Fig. 2D and 2E) suggest that S6K1/2 activity does not play any role in regulating the interaction between PDCD4 and eIF3F.

### PDCD4 and eIF3F regulate each other’s levels and Bcl-xL levels

To demonstrate that PDCD4 not only interacts with eIF3 but also regulates its levels, PDCD4 was depleted using siRNA, and the different subunits of eIF3 were analyzed by western blot. The levels of eIF3D, eIF3E, eIF3F (direct interaction partner of PDCD4), and eIF3G were significantly decreased when PDCD4 was depleted in U343 cells (Fig. 3A). Interestingly when eIF3F was depleted using siRNA, the levels of PDCD4 were diminished in U343 cells (Fig. 3B; left panel). Treatment of MG132 (a potent proteasomal inhibitor) prevented the degradation of PDCD4 in eIF3F-depleted U343 cells (Supple. Fig. 1), suggesting that depletion of eIF3F destabilizes PDCD4 at the proteasomal level.

**Figure 3:**
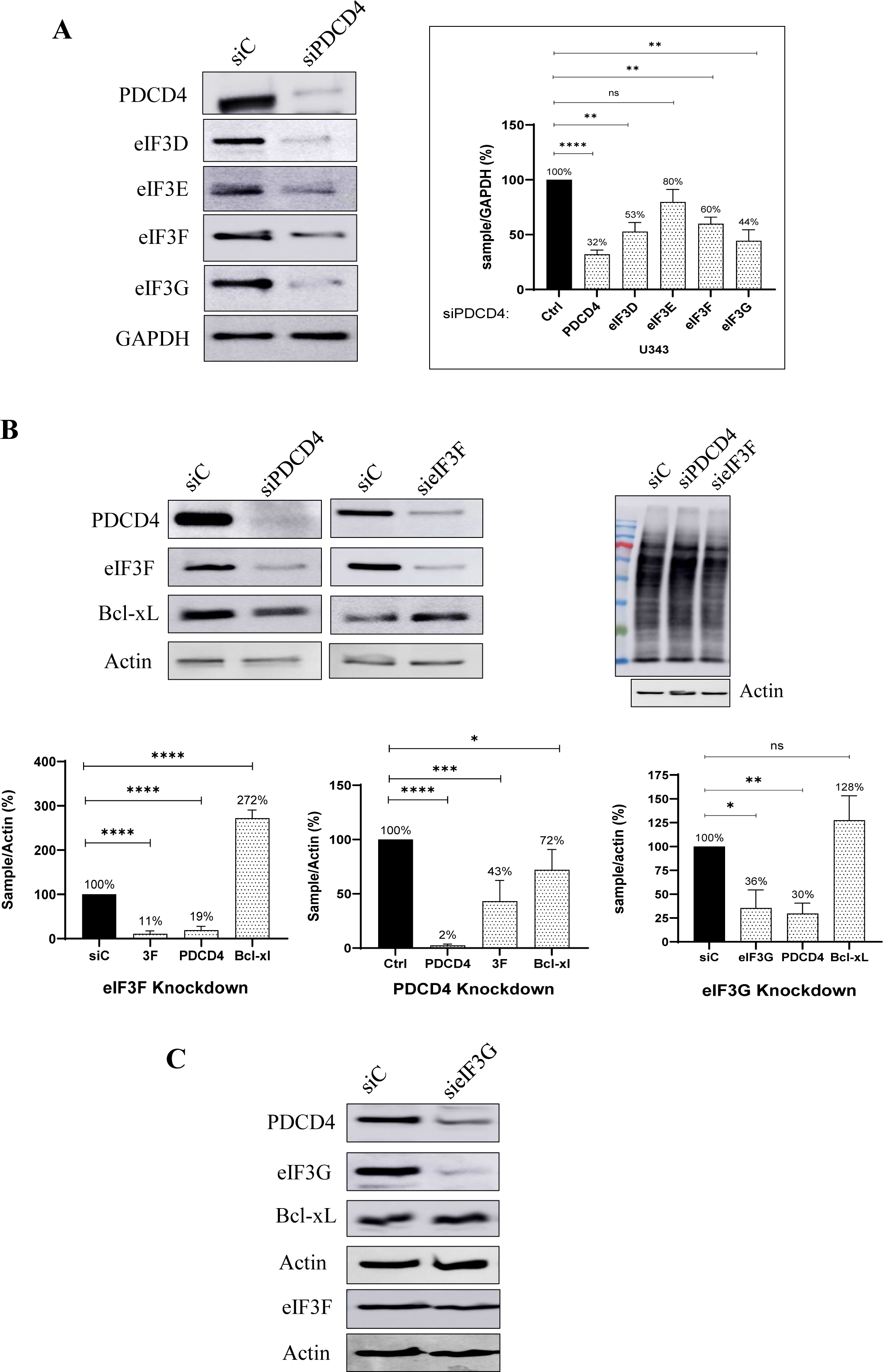
eIF3F depletion in U343 cells enhances the levels of Bcl-xL. Under PDCD4 depleted conditions, levels of eIF3D, eIF3E, eIF3F and eIF3G were reduced **(A)**. siRNA-mediated depletion of PDCD4 resulted in the decreased levels of eIF3F, and Bcl-xL (**B; left panel)**. When eIF3F was depleted, lower expression of PDCD4 was observed, however, the levels of Bcl-xL were robustly increased **(B; middle panel)**. Puromycin Incorporation Assay showed that PDCD4 and eIF3F depletion the overall translation was not affected **(B; right panel)**. In a control experiment, eIF3G was depleted from U343 cells and the levels of PDCD4, eIF3F, and Bcl-xL were determined. Upon eIF3G depletion, the levels of PDCD4 were reduced. However, the levels of eIF3F and Bcl-xL were not affected **(C)**. The graphs represent the quantified results of the western blot images. Significance is represented as nonsignificant – “ns”, p < 0.05 – “*”, p < 0.005 – “**, p < 0.001 – “***”, p < 0.0001 – “****”; (n=3, ± SEM).

Depletion of PDCD4 resulted in a significant reduction of Bcl-xL level (Fig. 3B). On the contrary, the levels of Bcl-xL greatly increased under eIF3F depletion condition in U343 cells (Fig. 3B). However, the global protein synthesis was not significantly affected by depletion of PDCD4 or eIF3F as revealed by the puromycin incorporation assay (Fig. 2B; right panel). This suggests that PDCD4 and eIF3F have a regulatory effect on each other and the levels of anti-apoptotic protein Bcl-xL (Fig. 3B). Depletion of eIF3G also decreased the levels of PDCD4. However, the levels of Bcl-xL were not affected by eIF3G depletion, confirming the exclusive impact of eIF3F on the regulation of Bcl-xL expression (Fig. 3C). These findings underscore the reciprocal regulatory influence between PDCD4 and eIF3F, suggesting their joint modulation of the anti-apoptotic protein Bcl-xL.

### PDCD4 acts via eIF3F in regulating the levels of Bcl-xL

In the preceding sections, we show the regulation of the anti-apoptotic protein Bcl-xL in PDCD4- and eIF3F-depleted U343 cells. Following this, we exogenously overexpressed PDCD4 or eIF3F in U343 cells. Overexpression of PDCD4 resulted in significant enhancement of Bcl-xL levels. However, the levels of eIF3F were not affected by PDCD4 overexpression (Fig. 4A and Fig. 1A). eIF3F overexpression resulted in a modest decrease in Bcl-xL levels without affecting the levels of PDCD4 (Fig. 4A). In order to check if PDCD4 and eIF3F co-regulate Bcl-xL expression, we conducted experiments involving the overexpression of eIF3F in PDCD4-depleted conditions and PDCD4 overexpression in eIF3F-depleted conditions. The levels of Bcl-xL were decreased under PDCD4 depletion under pcDNA3-FLAG control. (Fig. 4B; left panel). Overexpression of eIF3F under PDCD4 depletion condition further decreased the levels of Bcl-xL (Fig. 4B; left two panels and bottom left graph). Moreover, as expected, the levels of Bcl-xL were robustly increased under eIF3F depletion under control conditions. However, overexpression of PDCD4 under the eIF3F depletion condition resulted in a significant overall decrease in Bcl-xL levels (Fig. 4B; right two panels and bottom right graph). These results suggest that PDCD4 and eIF3F protein levels are critical in the regulation of Bcl-xL and PDCD4 exerts its effect in regulating Bcl-xL via eIF3F.

**Figure 4:**
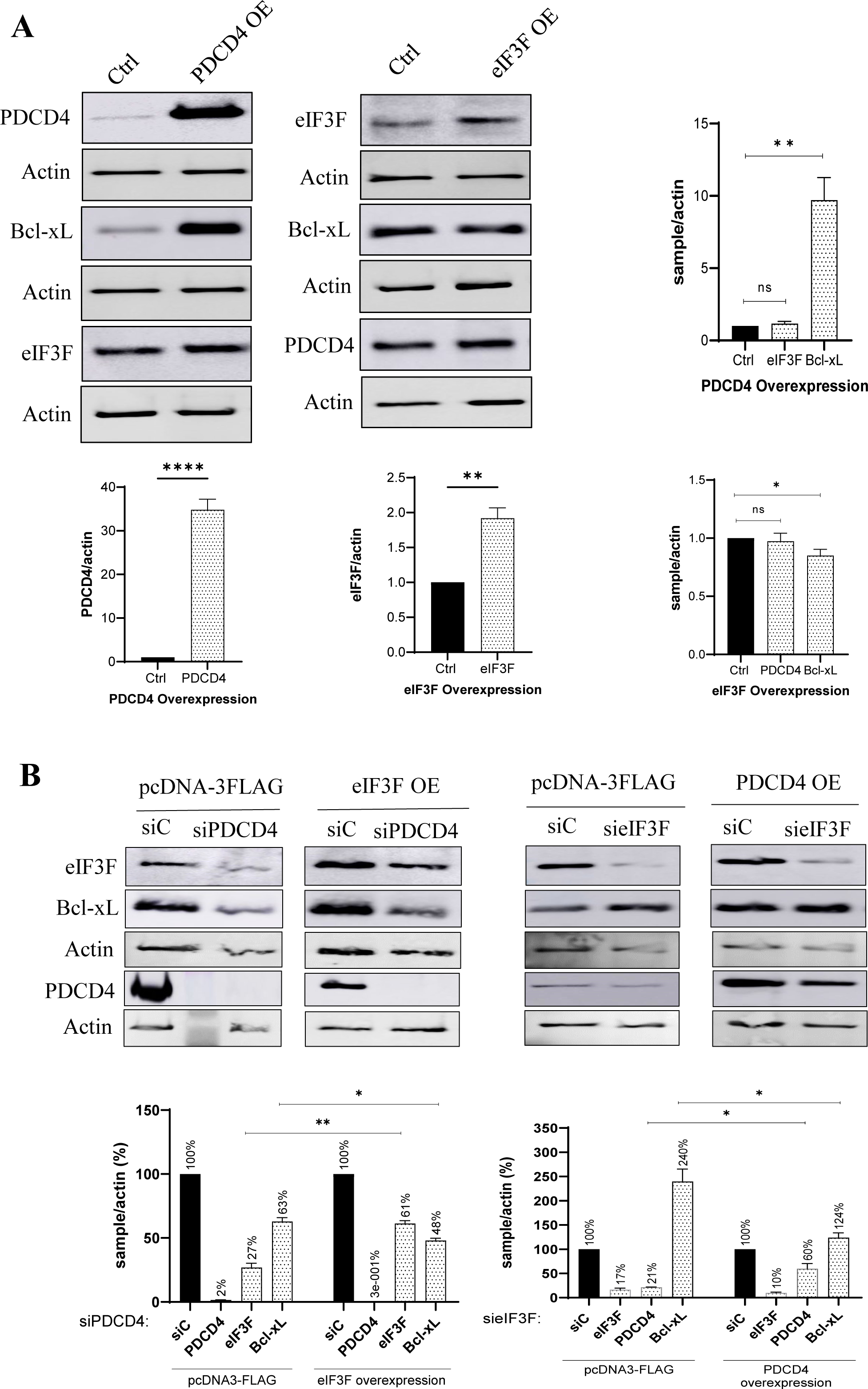
PDCD4 acts via eIF3F in regulating the levels of Bcl-xL. Overexpression of PDCD4 resulted in a robust enhancement of Bcl-xL levels without affecting the levels of eIF3F **(A)**. eIF3F overexpression showed a significant (but not robust) decrease in Bcl-xL without affecting the levels of PDCD4 **(A)**. We exogenously overexpressed eIF3F in PDCD4 depletion conditions and PDCD4 in eIF3F depletion conditions. As expected, the levels of Bcl-xL were decreased under the PDCD4 depletion + pcDNA-3FLAG (control) condition. Overexpression of eIF3F under PDCD4 depletion condition further decreased the levels of Bcl-xL (modest but significant decrease; compared to PDCD4 depletion + pcDNA-3FLAG (control) condition) **(B; left two panels and bottom left graph)**. Moreover, as expected, the levels of Bcl-xL were robustly increased under the eIF3F depletion + pcDNA-3FLAG (control) condition. However, overexpression of PDCD4 under eIF3F depletion condition resulted in a significant overall decrease in Bcl-xL levels (compared to eIF3F depletion + pcDNA-3FLAG (control) condition) **(B; right two panels and bottom right graph**). This data suggests that PDCD4 exerts its effect in regulating Bcl-xL via eIF3F. The graphs represent the quantified results of the western blot images. Significance is represented as nonsignificant – “ns”, p < 0.05 – “*”, p < 0.005 – “**, p < 0.001 – “***”, p < 0.0001 – “****”; (n=3, ± SEM).

### Depletion of eIF3F greatly enhances the translation of Bcl-xL mRNA

The observation of RNA-independent interaction between PDCD4 and eIF3F (Fig. 1) raises the question of their individual associations with Bcl-xL RNA. To this end, we perform endo-RNA-IP using PDCD4 and eIF3F antibodies. We performed RT-qPCR analysis to analyze the interaction of Bcl-xL mRNA with PDCD4 and eIF3F. Compared to negative control (FBS), the levels of Bcl-xL mRNA were a higher in PDCD4 and eIF3F endo-IP (Fig. 5A). Bcl-xL mRNA was enriched in these endo-IPs compared to input (lysate) (Fig. 5A). This suggest that both PDCD4 and eIF3F bind to the Bcl-xL RNA (Fig. 5A). Therefore, we wanted to investigate if the levels of PDCD4 and eIF3F affect each other’s ability to interact with Bcl-xL mRNA. In this regard, we depleted PDCD4 or eIF3F from U343 cells using shRNAs. Subsequently, we performed endo-RNA-IP as mentioned above. Under the PDCD4 depletion condition, eIF3F efficiently interacted with Bcl-xL mRNA (Fig. 5B). Likewise, under the eIF3F depletion condition, PDCD4 efficiently interacted with Bcl-xL mRNA (Fig. 5B). There was no significant difference in the interaction between PDCD4 and Bcl-xl mRNA or eIF3F and Bcl-xL mRNA (shC vs. shPDCD4/sheIF3F) (Fig. 5B). This suggests that PDCD4 and eIF3F neither enhance nor inhibit each other’s interaction with Bcl-xL mRNA. Both these protein factors bind independently to Bcl-xL mRNA and they do not compete or cooperate to bind to Bcl-xL mRNA (Fig. 5A, 5B).

**Figure 5:**
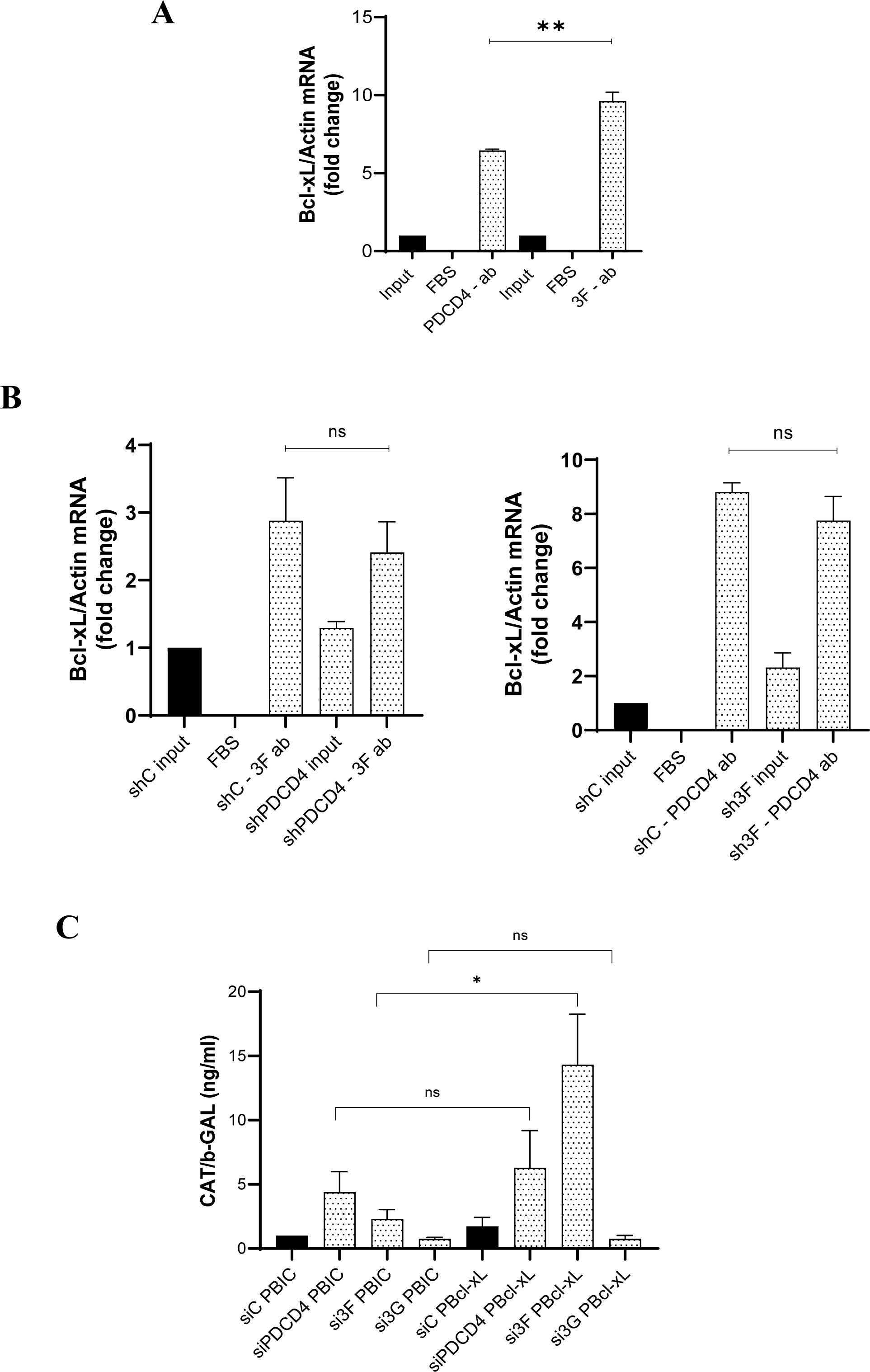
PDCD4 and eIF3F interact with the Bcl-xL mRNA and eIF3F depletion enhances Bcl-xL IRES activity. *Endo RNA IP* results show that PDCD4 and eIF3F interact with Bcl-xL RNA. There was a significant increase observed between PDCD4 and eIF3F association with Bcl-xL RNA. FBS was used as a negative control and all samples were normalized internally with actin mRNA. Input samples were used for the normalization of PDCD4-FBS, PDCD4-ab, eIF3F-FBS, and eIF3F-ab samples **(A)**. shRNA-mediated depletion of PDCD4 did not enhance the interaction of eIF3F with Bcl-xL mRNA **(B; left panel)**. Also, shRNA-mediated depletion of eIF3F did not enhance the interaction of PDCD4 with Bcl-xL mRNA **(B; right panel)**. These results suggest that PDCD4 and eIF3F interact with Bcl-xL mRNA independently and they do not compete or cooperate to bind to Bcl-xL mRNA. Bicistronic reporter assay showed that depletion of PDCD4 did not affect Bcl-xL IRES activity, whereas the depletion of eIF3F robustly and significantly enhanced Bcl-xL IRES activity. However, as expected, Bcl-xL IRES activity was not affected by the depletion of eIF3G (control) **(C)**. Significance is represented as nonsignificant – “ns”, p < 0.05 – “*”, p < 0.005 – “**, p < 0.001 – “***”, p < 0.0001 – “****”; (± SEM, n=3).

As the 5’ UTR of Bcl-xL mRNA is known to harbor IRES element [38], we investigated the ability of PDCD4 and eIF3F to regulate IRES-mediated translation of Bcl-xL mRNA in U343 cells. We depleted eIF3G (negative control), eIF3F, or PDCD4 from U343 cells. Subsequently, cells were transfected with plasmids containing the bicistronic reporters, β-galactosidase (β-gal), and chloramphenicol acetyltransferase (CAT), including Bcl-xL IRES element in between them (pBIC as control and pBIC-Bcl-xL IRES including the IRES element). The IRES activity was measured by taking a ratio of CAT/β-gal levels. As expected, Bcl-xL IRES activity was not affected by eIF3G depletion (Fig. 5C). Also, there was a non-significant change in IRES activity when PDCD4 was depleted. However, depletion of eIF3F from U343 cells resulted in a significantly increased level of r Bcl-xL IRES activity (Fig. 5C). This suggests that eIF3F normally inhibits IRES-mediated translation of Bcl-xL, however, the depletion of eIF3F from GBM cells enhances the IRES-mediated translation of Bcl-xL.

In order to check if the translation of endogenous Bcl-xL mRNA is regulated by eIF3F or PDCD4, we performed a polysome profiling assay. To this end, we efficiently depleted PDCD4 or eIF3F from U343 cells using shRNAs (Fig. 6A). shRNA-mediated depletion of PDCD4 did not significantly affect the steady-state mRNA levels of eIF3F and Bcl-xL mRNAs (Fig. 6B; left panel). Likewise, shRNA-mediated depletion of eIF3F did not significantly affect the steady-state mRNA levels of PDCD4 and Bcl-xL (Fig. 6B; right panel). We separated monosomes and polysomes in sucrose density gradient using ultracentrifugation from the control, PDCD4- or eIF3F-depleted U343 cells (Fig. 6C). The fraction collected were subjected to RT-qPCR analysis using ΔΔCt method relative to actin mRNA levels to determine the % distribution of Bcl-xL, eIF3F, or PDCD4 mRNA over 10 fractions of the polysome profile (Suppl. Fig. 2). In PDCD4-depleted U343 cells the translation of mRNAs encoding eIF3F and Bcl-xL did not change significantly (Suppl. Fig. 2). Additionally, under eIF3F-depletion condition, the translation of PDCD4 mRNA did not change (Supple. Fig. 2). However, the translation of Bcl-xL mRNA was greatly increased in eIF3F-depleted U343 cells (Supple. Fig. 2). Also, by taking ratios of Bcl-xL mRNA in polysome fractions (5-10) over Bcl-xL mRNA in monosome fraction (1-4), we confirmed that the translation of Bcl-xL mRNA was not significantly affected by the depletion of PDCD4 (Fig. 6C; right panel). However, the translation of Bcl-xL mRNA was robustly and significantly increased in eIF3F-depleted U343 cells (Fig. 6C; right panel).

**Figure 6:**
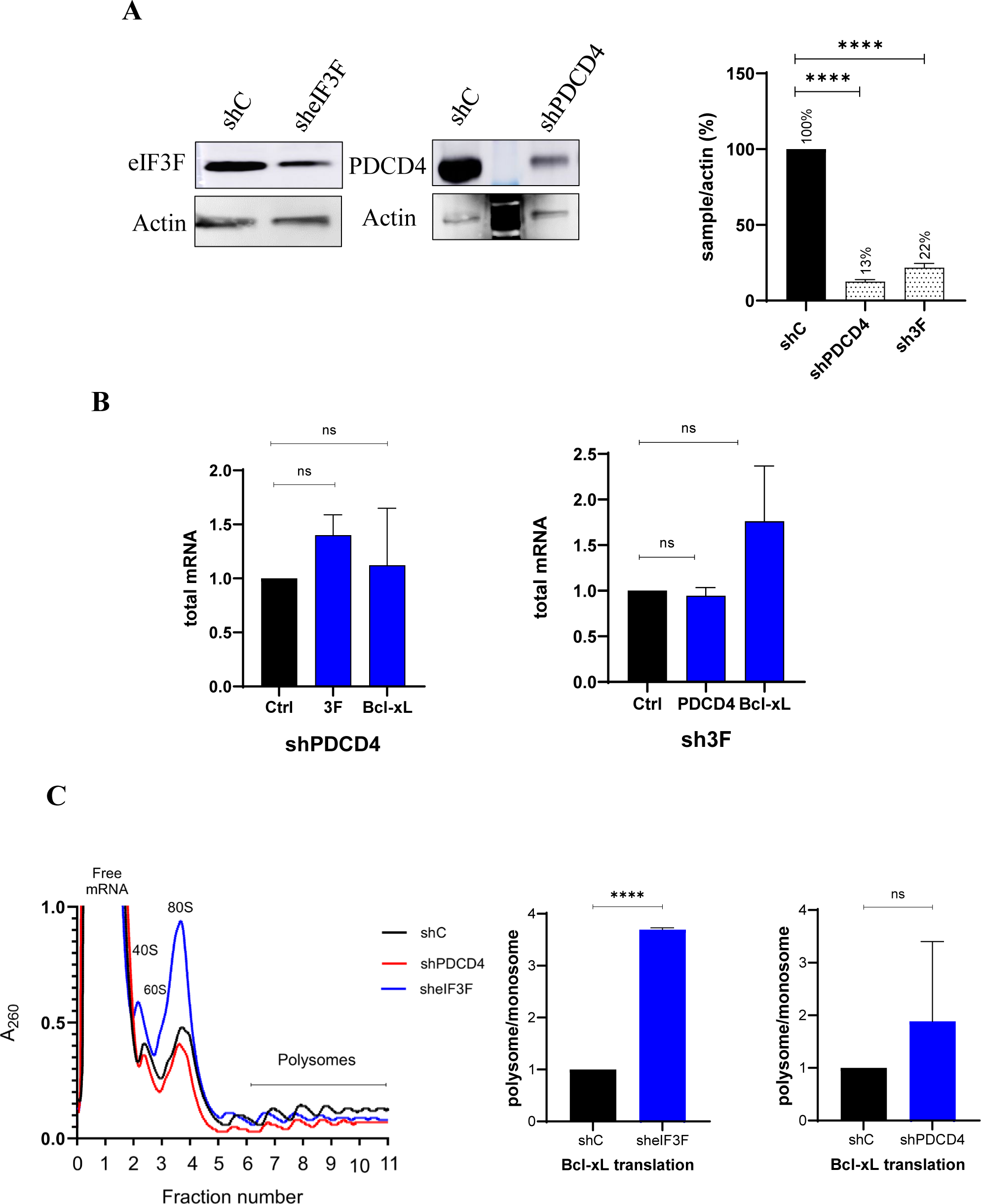
eIF3F depletion enhanced the translation of Bcl-xL mRNA. shRNA-mediated depletion of PDCD4 and eIF3F was confirmed by western blot analysis **(A).** Total mRNA levels of eIF3F and Bcl-xL were analyzed in cells transduced with shPDCD4. Total mRNA levels of PDCD4 and Bcl-xL were analyzed in cells transduced with sheIF3F **(B)**. Polysome profile of shC, shPDCD4, and sheIF3F showing the monosomes and polysomes levels across multiple fractions **(C; left panel)**. Polysome to Monosome ratios show that Bcl-xL mRNA translation was robustly and significantly increased in eIF3F depletion condition (C; middle panel). Depletion of PDCD4 did not significantly affect the mRNA translation of Bcl-xL **(C; right panel)**. Significance is represented as nonsignificant – “ns”, p < 0.05 – “*”, p < 0.005 – “**, p < 0.001 – “***”, p < 0.0001 – “****”; (± SEM, n=3).

### eIF3F-PDCD4 dependent regulation of Bcl-xL is specific to GBM cells

We depleted PDCD4 or eIF3F in three different glioblastoma cell lines, one of which is a primary brain tumor stem cell (BTSC) line. For comparison, we depleted PDCD4 or eIF3F in human lung fibroblasts. Subsequently, we performed western blot analysis for PDCD4, eIF3F, and Bcl-xL in all 4 cell lines. Depletion of PDCD4 in glioblastoma cells U343 (Fig. 3B), U373 (Fig. 7A), and BT73MR (Fig. 7B) resulted in a significant reduction of Bcl-xL level (Fig. 3B). On the contrary, the levels of Bcl-xL greatly increased under eIF3F depletion condition in U343 (Fig. 3B) U373 (Fig. 7A), and BT73MR (Fig. 7B) cells. When PDCD4 was depleted in human lung fibroblast (WI-38) cells, the levels of eIF3F and Bcl-xL were not affected (Fig. 7C). However, there was a modest increase in PDCD4 and Bcl-xL levels in eIF3F-depleted WI-38 cells (Fig. 7C). This result indicates that the PDCD4 and eIF3F mediated regulation of Bcl-xL occurs predominantly in glioblastoma cells.

**Figure 7:**
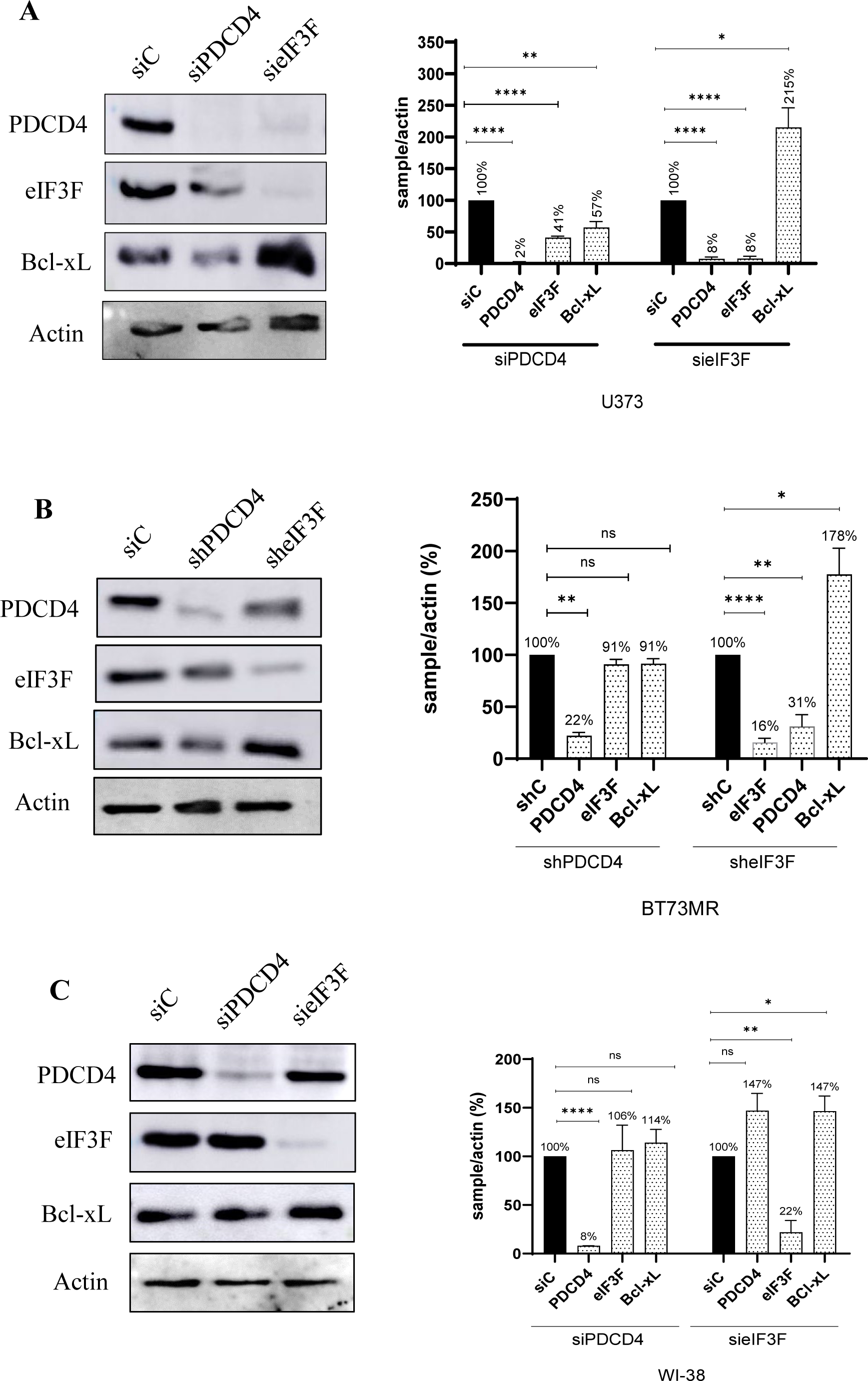
eIF3F-PDCD4 dependent mechanism is implicated in glioblastoma cells. Like in U343 cells, eIF3F depletion in U373 cells robustly enhanced the levels of Bcl-xL. Also, the depletion of PDCD4 in U373 cells resulted in a significant decrease in Bcl-xL levels **(A).** Depletion of eIF3F, and PDCD4 decreased the levels of PDCD4 and eIF3F, respectively **(A)**. Similarly, eIF3F depletion in BT73MR robustly enhanced the levels of Bcl-xL. Also, the depletion of PDCD4 in brain tumor stem cells (BT73MR) resulted in a significant decrease in Bcl-xL levels **(B).** Depletion of eIF3F, and PDCD4 decreased the levels of PDCD4 and eIF3F, respectively **(B)**. WI-38 did not show decreased levels of eIF3F in PDCD4 depletion or vice versa, which is strikingly different from GBM cells. The levels of Bcl-xL in eIF3F depletion were significantly high, however, not as robust compared to GBM cells **(C).** WI-38 cells did not show decreased levels of eIF3F in PDCD4 depletion or vice versa, which is remarkably different from GBM cells. Also, the levels of Bcl-xL in eIF3F or PDCD4 depletion conditions were not significantly affected **(D)**. Significance is represented as nonsignificant – “ns”, p < 0.05 – “*”, p < 0.005 – “**, p < 0.001 – “***”, p < 0.0001 – “****”; (n=3, ± SEM).

## Discussion

Regulation of mRNA translation plays a critical role in cellular homeostasis and disease [39, 40]. During cellular stress, it is observed that the mRNA profile is not consistent with the proteome profile of the cell [41, 42]. This suggests that when it comes to translation not all transcripts are treated equally [42, 43]. This underlines the importance of non-canonical mechanisms of translation regulation under cellular stress conditions. IRES-mediated translation initiation is one of the key modes of non-canonical translation initiation that regulates the translation of a subset of mRNA [4, 7, 10]. Largely, the translation of stress-related mRNAs (such as pro- and anti-apoptotic proteins) is regulated by IRES-mediated translation initiation [44, 45].

IRES transacting factors (ITAFs) play an important role in regulating IRES-mediated translation initiation [10]. Investigating these ITAFs for their role in IRES-mediated translation initiation is critical as they could be the immediate option for the cells to survive under stress conditions without de novo mRNA synthesis.

PDCD4 is a well-known tumor suppressor protein, which also binds to eIF4A and PABP to regulate canonical translation initiation [20]. Our work established PDCD4 as an ITAF that regulates IRES-mediated translation of XIAP and Bcl-xL mRNAs [19, 23]. eIF3F is a subunit of the large initiation factor, eIF3, that plays a critical role in canonical and non-canonical translation initiation [26].

Both PDCD4 and eIF3 are targets of the mTOR pathway [46] and are phosphorylated by S6K1 and/or 2, a downstream target of mTOR [46]. PDCD4 phosphorylation promotes PDCD4 degradation via ubiquitination while eIF3 phosphorylation enhances eIF3 activity [32, 46]. One of the known common target mRNAs for PDCD4 and eIF3 is XIAP mRNA [23, 33]. PDCD4 and eIF3 directly interact with XIAP IRES RNA and have opposing effects on XIAP mRNA translation initiation [23, 33]. PDCD4 is also shown to directly interact with Bcl-xL IRES RNA and to inhibit IRES-mediated translation of Bcl-xL. Moreover, PDCD4 was shown to interact with eIF3H and to enhance metastasis of lung adenocarcinoma [47]. However, the mRNA translation mechanism of PDCD4-eIF3 interaction remained largely unexplored. Hence, this study investigates the interaction between PDCD4 and eIF3 as this could be a possible mechanism of Bcl-xL mRNA translation regulation.

The experiments demonstrate a specific RNA-independent and direct interaction between PDCD4 and the eIF3 complex. eIF3 is a large 13-subunit complex of which subunits, eIF3B, eIF3D, eIF3E, eIF3G, and eIF3F subunits were shown to interact with PDCD4 in the co-IP performed (Fig. 1). Based on the above results, it was hypothesized that the co-immunoprecipitation of these eIF3 subunits could be a result of the following three types of interactions. The interaction between PDCD4 and eIF3 has the potential to impede eIF3 association with mRNA or other cellular proteins, thereby exerting inhibitory effects on translation. eIF3 may comprise a core complex composed of specific subunits, with the remaining subunits potentially participating exclusively in regulatory mechanisms. PDCD4 could interact with a subcomplex encompassing eIF3B, eIF3D, eIF3E, eIF3G, and eIF3F subunits, leading to the inhibition of translation. Additionally, PDCD4 might establish direct interactions with individual subunits, as demonstrated for eIF3F in this study. Such interactions could hinder the subunit’s association with other eIF3 components, thereby preventing the formation of a functional complex, or impeding its autonomous activity.

eIF3F has also been earlier reported to interact with S6K1 and was demonstrated by an *in vitro* experiment [32]. S6K1 binds to eIF3F to phosphorylate eIF3 subunits [32]. However, the *in cellulo* experiments suggested that both the S6K1 and 2 have a role in eIF3 phosphorylation [32]. PDCD4 interacting with eIF3F can interfere with eIF3F and S6K1/2 association to prevent phosphorylation of the eIF3 complex. Similarly, PDCD4 binding to eIF3B and eIF3E may also affect interaction with eIF4G and inhibit ribosome recruitment to the mRNA. These canonical and specialized functions of the eIF3 subunits can be affected by their interaction with PDCD4. Besides eIF3, PDCD4 is known to interact with PABP and eIF4A to suppress their role in translation initiation [20, 21]. Therefore, PDCD4-eIF3 interaction may have an inhibitory effect on translation initiation.

Phosphorylation of proteins can alter the charge, structure, and binding affinity of the proteins [48–50]. Interestingly, both PDCD4 and eIF3 are phosphorylated by S6K1 and/or 2 [32] which could affect their interaction. However, our data indicates that S6K1 and/or 2 activities did not affect the PDCD4-eIF3F interaction. Treatment with serum-free media could trigger multiple effects besides S6K1 and 2 inhibitions. Therefore, it is important to perform similar co-IP experiments using chemical inhibitors that specifically target S6K1 and 2 activities. Pharmacological inhibition of mTOR/S6K1/2 axis did not affect the PDCD4-eIF3F interaction (Fig. 2E). This result of eIF3F and PDCD4 interaction during serum-starvation condition or pharmacological inhibition of S6K1/2 may not be extrapolated to all the eIF3 subunits, as the interaction between each of the eIF3 subunit and S6Ks may vary. According to our results, S6K1 and S6K2 or inhibition of them did not affect PDCD4-eIF3F interactions.

Not only do PDCD4 and eIF3/eIF3F interact with each other, but we also show that they regulate each other at the protein level. When PDCD4 was depleted, we observed a decrease in subunits of eIF3 such as eIF3D, eIF3E, eIF3F, and eIF3G. This regulation seems to be reciprocal as when we depleted eIF3F or eIF3G, we observed a reduction in PDCD4 levels (Fig. 3). These observations suggest that eIF3 (specifically eIF3F & eIF3G) and PDCD4 have a stabilizing effect on each other. Moreover, under conditions of cellular stress, specific proteins undergo targeted degradation via the proteasomal pathway as part of regulatory mechanisms. PDCD4 is phosphorylated by S6K2 and is targeted for proteasomal degradation [23]. Our data clearly shows that post eIF3F depletion, the levels of PDCD4 were recovered when treated with a synthetic proteasomal degradation inhibitor MG132. Therefore, it is very likely that PDCD4 and eIF3F work in synergy to regulate the IRES-mediated translation of anti-apoptotic protein Bcl-xL.

Interestingly, when we depleted PDCD4, we noticed a decrease in Bcl-xL protein levels, whereas we found a significant two-fold increase in Bcl-xL levels when eIF3F was depleted (Fig. 3). These findings demonstrate the critical role of eIF3F in the regulation of Bcl-xL protein. As depletion of eIF3F or PDCD4 affects each other’s protein level, one can expect that Bcl-xL levels would increase in both eIF3F- and PDCD4-depleted cells. However, it is important to note that depletion of eIF3F results in an extremely low level of PDCD4. In contrast, under PDCD4 depletion conditions, the decrease in eIF3F level was only ∼ 57% (Fig. 3). It is also possible that the molecular stoichiometry between PDCD4 and eIF3F influences the IRES-mediated translation of Bcl-xL. Therefore, with the establishment of a direct interaction between PDCD4 and eIF3F, we propose that PDCD4 operates through eIF3F to upregulate Bcl-xL. When PDCD4 was over-expressed in eIF3F-depleted cells, the levels of Bcl-xL did not show a significant increase as compared to the Bcl-xL levels in control (pcDNA3-FLAG)eIF3F-depleted cells (Fig. 4). Upon overexpression of eIF3F in conditions where PDCD4 was depleted, there was a moderate increase in the levels of Bcl-xL. This strongly suggests that the PDCD4 and eIF3F interaction influences the regulation of Bcl-xL. Findings from the overexpression experiment indicate that PDCD4 functions through eIF3F to regulate Bcl-xL at the protein level. However, additional validation is required to confirm the regulatory mechanisms at the translation level and the modulation of IRES-mediated translation.

By performing RNA-IP experiments, we showed that eIF3F and PDCD4 interact with Bcl-xL mRNA directly and they do not compete or cooperate for binding to Bcl-xL mRNA (Fig. 5). We further analyzed the effect of PDCD4 and eIF3F on IRES-mediated translation initiation of Bcl-xL. The results of the bicistronic reporter assay indicated that the levels of IRES activity were higher in the eIF3F knockdown condition compared to PDCD4 depletion (Fig. 5C). By polysome profiling, we have demonstrated that eIF3F regulates Bcl-xL levels at the mRNA levels. There is a robust increase in Bcl-xL mRNA translation when eIF3F is depleted, however, there was no statistically significant difference in Bcl-xL mRNA translation when PDCD4 was depleted. Puromycin incorporation assay reported that the was no increase in overall protein synthesis in PDCD4 and eIF3F depletion (Fig. 3C). Hence, the elevation in Bcl-xL levels did not exhibit proportionality to the overall increase in translation. In eIF3F depletion, we did not detect a decrease in PDCD4 mRNA translation as we have seen at the protein level. This could be explained by the regulation of PDCD4 at the proteasomal level. Together, all the results suggest that PDCD4 and eIF3F interaction is crucial for the regulation and expression of Bcl-xL. Furthermore, it is possible that PDCD4 exerts a negative regulatory influence on Bcl-xL expression by interacting with eIF3F, and in there, their stoichiometry would be important.

The overall study aims at deciphering the mechanism of regulation of mRNA translation of an anti-apoptotic protein, Bcl-xL, by PDCD4 and eIF3F. In conclusion, this study suggests that PDCD4 and eIF3F interact with each other directly in an RNA-independent manner. At the protein level, PDCD4 is regulated at the proteasomal level. PDCD4 and eIF3F negatively regulate each other while regulating the anti-apoptotic protein Bcl-xL. This study has revealed that Bcl-xL expression was mediated by PDCD4 through eIF3F, eIF3F being the key regulator. We also demonstrated that eIF3F depletion activates IRES-mediated translation initiation of Bcl-xL. Moreover, by depleting tumor suppressors PDCD4 or eIF3F in at least three glioblastoma cell lines (including brain tumor stem cells) and non-cancer human lung fibroblasts, we show that PDCD4-eIF3F-dependent regulation of Bcl-xL is likely specific to glioblastoma cells. Thereby, this mechanism may play an important role in gliomagenesis and glioblastoma pathophysiology.

## Acknowledgments

This work was funded by a Natural Sciences and Engineering Research Council of Canada-Discovery Grant (RGPIN-2017-05463), the Canada Foundation for Innovation-John R. Evans Leaders Fund (35017), the Campus Alberta Innovates Program and the Alberta Ministry of Economic Development and Trade. We thank Dr. Tithi Ghosh for critically reading this manuscript.

## Author Contributions

Conceptualization, NT, MH, VH, DKS; Methodology, VH, DKS; HP, PN, JL; Analysis,VH, DKS, PN; Investigation, VH, DKS; HP, PN, JL; Resources, NT, MH; Writing – Original Draft, VH, DKS, NT; Writing – Review and Editing, NT, MH; Supervision, NT, MH; Funding Acquisition, NT

## Declaration of Interests

The authors declare no competing interests.

## Notes

### Competing Interest Statement

The authors have declared no competing interest.

