## Supplemental Information for "Eukaryotic initiation factor 3F (eIF3F) regulates the IRES-mediated translation of Bcl-xL via its interaction with programmed cell death 4 (PDCD4) protein"

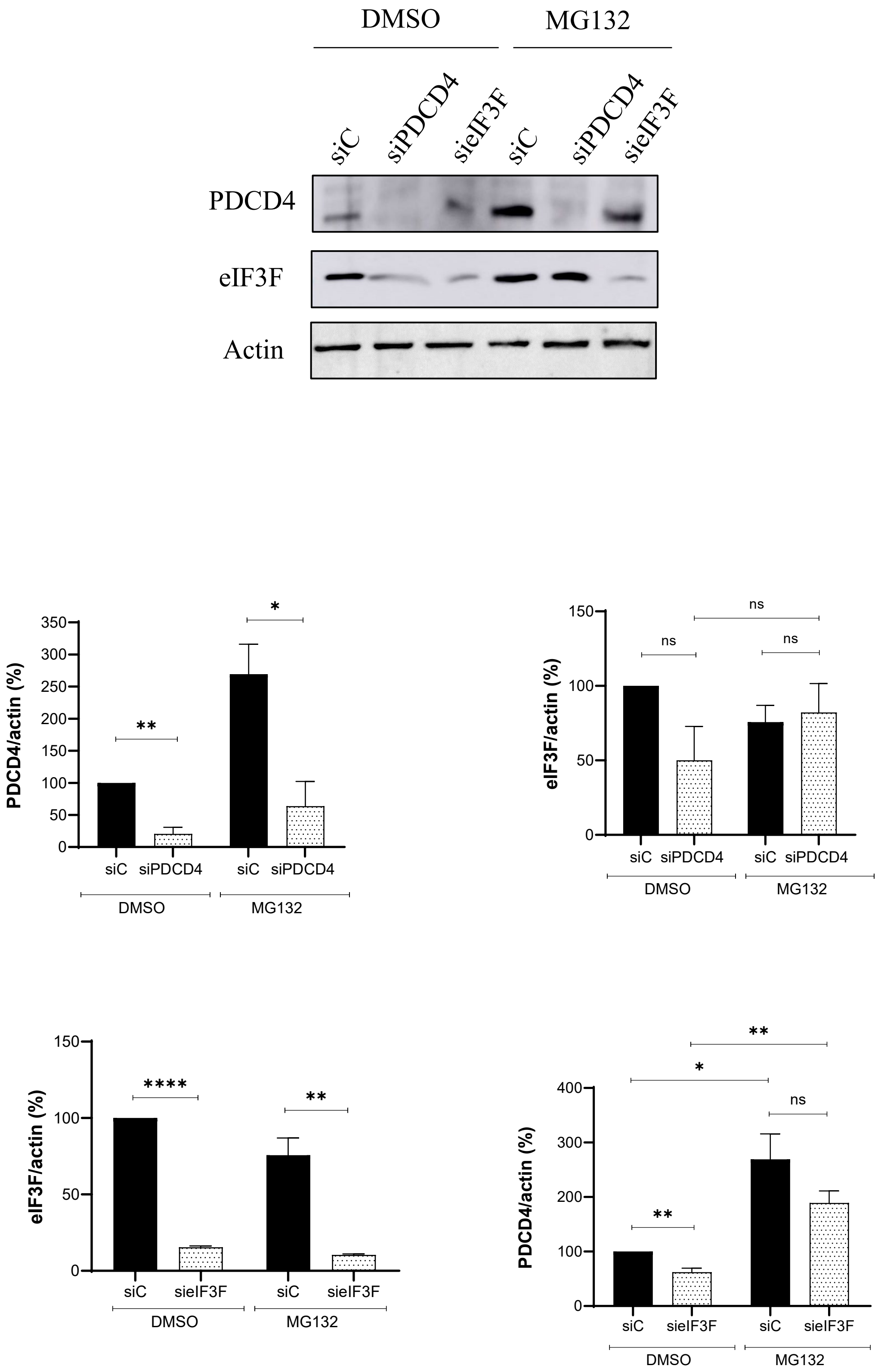


Supplementary Figure 1

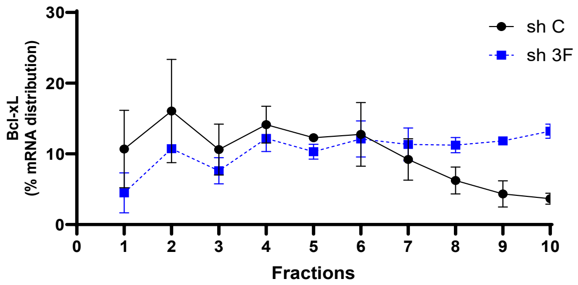


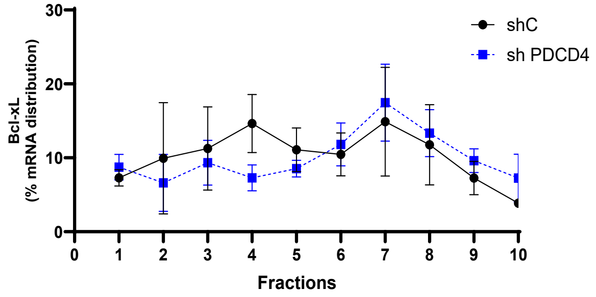


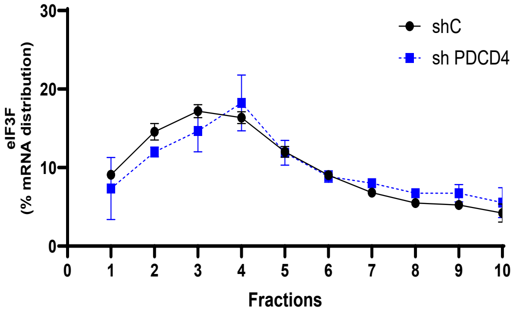


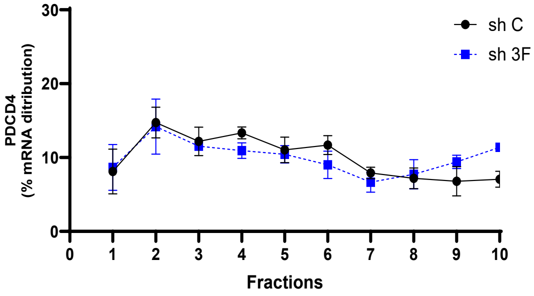


Supplementary Figure 2

Supplemental table 1

List of constructs, application of the constructs and their source.

| Sl. No | Construct | Application | Source | Reference |
|---|---|---|---|---|
| 1 | 2G-T.f | Bacterial expression of GST-eIF3F | Dr. Jamie H. D. Cate, University of California, Berkeley. | (Sun et al., 2011) |
| 2 | His-PDCD4 | Bacterial expression of His-PDCD4 | Dr. Martin Holcik, CHEO Research Institute, University of Ottawa | (Liwak et al., 2012) |
| 3 | pcDNA3-PDCD4-FLAG | Mammalian expression of FLAG-PDCD4 | | |
| 4 | GST-PDCD4 | Bacterial expression of GST-PDCD4 | | |
| 5 | pcDNA3-FLAG | Mammalian expression of FLAG-tag | | |
| 6 | HA-eIF3F | Mammalian expression of HA-eIF3F | Dr. Nahum Sonenberg McGill University | (Martineau, Y., et al, MCB, 2014) |
| 7 | βGAL-CAT (PBIC) | Bicistronic Reporter Assay | Dr. Martin Holcik, CHEO Research Institute, University of Ottawa | (Liwak et al., 2012) |
| 8 | βGAL – Bcl-xL – IRES – CAT (pBcl-xL) | Bicistronic Reporter Assay | Sigma Millipore – Target sequence: CAGTCACAGATTGCACTCAAT | |
| 9 | sh87 (eIF3F) (# TRCN0000073987) | Polysome profiling | | |
| 10 | Sh79 (PDCD4) (#TRCN0000059079) | Polysome profiling | Sigma Millipore – Target sequence: GCGGTTTGTAGAAGAATGTTT | - |
| 11 | psPax2 (packaging plasmid) | Lentivirus production | Addgene **-**Plasmid #12260 | Gift from Didier Trono |
| 12 | pMD2.G (envelope plasmid) | Lentivirus production | Addgene – Plasmid #12259 | Gift from Didier Trono |

**Supplemental Table 2:** The list of antibodies used and their source.

| Antibody | Source |
|---|---|
| PDCD4 | Rockland (#600-401-965), Proteintech (#12587-1-AP), Abcam (#ab80590) |
| eIF3 subunits: eIF3B, eIF3D, eIF3E, eIF3F, eIF3G, eIF3H | **Abcam**<br>eIF3B (#ab133601), eIF3D (#ab155419), eIF3E (#ab36766), eIF3F (#ab74568), eIF3G (#ab192601), eIF3H (#ab228536) |
| S6 | Cell signaling technology (#2217) |
| pS6 | Cell signaling technology (#4858S) |
| Bcl-xL (polyclonal and monoclonal) | Cell signaling technology (#2762S, #2764S) |
| XIAP (polyclonal and monoclonal) | Cell signaling technology (#2042S), Proteintech (#66800-1-Ig) |
| Actin | Bio-Rad (#12004163) |
| Anti-rabbit (secondary) | Abcam (#ab97051) |
| Anti-mouse (secondary) | Abcam (#ab6728) |
| Conformation-specific anti-rabbit | Cell signaling technology (#5127S) |
